# Effectiveness of a mobile antiretroviral pharmacy and HIV care intervention on the continuum of HIV care in rural Uganda

**DOI:** 10.1101/533950

**Authors:** Francis Bajunirwe, Nicholas Ayebazibwe, Edgar Mulogo, Maria Eng, Janet McGrath, David Kaawa-Mafigiri, Peter Mugyenyi, Ajay K. Sethi

## Abstract

**Introduction:** Adherence to antiretroviral therapy (ART) is critical in order to achieve viral suppression, one of three UNAIDS targets set for achievement before 2020. One of the main barriers to adherence is the long distance between patient residences and healthcare facilities. We designed an intervention, Mobile Antiretroviral Therapy and HIV care (MAP-HC) in rural southwestern Uganda aimed to reduce travel distance and hypothesized that MAP-HC would improve ART adherence and rates of viral load suppression.

**Methods:** The study was conducted at two sites, Kitagata and Itojo Hospitals, and these are public health facilities located in rural southwestern Uganda. Patients who lived >5km from the hospital were provided the option to participate. For each hospital, we identified 4 health centres in the catchment area to serve as site for the mobile pharmacy. Each site was visited once a month to provide ART refills, adherence counseling and treatment of other illnesses. We measured patient waiting time, adherence and viral load suppression before and after the intervention.

**Results:** We conducted baseline assessment among 292 patients at the two hospitals. The mean waiting time at Kitagata Hospital changed from 4.48 hours before the intervention but increased to 4.76 hours after the intervention (p=0.13). The proportion of patients who missed an ART dose in the last 30 days dropped from 20% at baseline to 8.5% at 12 months after the intervention (p=0.009). The proportion of patients with detectable viral load from 19.9% to 7.4% after the intervention (p=0.001).

**Conclusions:** Our study has showed that a mobile pharmacy intervention in rural Uganda is feasible and resulted in improvement in adherence and viral load suppression. Although it did not reduce patient waiting time at the clinic, we recommend a scale-up of this intervention in rural areas where patients face challenges of transportation to the clinic.

## Introduction

Adherence to antiretroviral therapy (ART) is critical in order to achieve viral suppression [1] or the third 90, one of three UNAIDS targets set for achievement before 2020. In order to achieve viral load suppression, patients should take at least 95% of their medications [2–4]. Although adherence levels in sub Saharan Africa are on average higher than estimates in North America [5], patients in countries such as Uganda face structural barriers to achieve high level adherence.

One of the main barriers to adherence is the long distance between one’s place of residence and healthcare facilities. [6, 7] Specifically, patients seeking HIV care and treatment travel longer distances compared to their HIV negative counterparts. [8] Several studies have examined geographical factors as barriers and systematic analysis has shown that travel distance is a barrier across the continuum of HIV care from testing to treatment and retention in care [9]. Patients narrate how they struggle to raise the monthly transportation fee to the clinics to collect their medicines [10, 11]. Some patients devised innovative ways such as pooling resources in order to secure their monthly pills, with some patients making two-day arduous journeys to an HIV clinic. Long waiting times because clinics are crowded [12] cause patients to view their monthly visits as burdensome, competing with time needed to tend to gardens and other income generating activities.

The World Health Organization now recommends the testing and initiation of ART for all persons infected with HIV regardless of CD4 count or HIV stage [13], referred to the “test and treat approach.” Uganda, along with other HIV-affected countries across the globe, is now implementing this strategy. The expected result is that HIV treatment clinics will experience rising numbers of patients needing treatment, worsening crowding and waiting times at these facilities. Such unintended effects of ART expansion are barriers to maintaining HIV care continuation, leading to poor retention and or non-adherence.

There is now a need for interventions that not only reduce distance to HIV treatment clinics, but also have potential to decongest them. These may result in better adherence to clinic visits, medications and improve viral load suppression. There is very limited interventions of this nature that have been tested in resource-limited settings. The few experiences with decentralized and community-based models in Africa have been successful [14, 15] including more recently from Nigeria where a community-based model increased adherence and retention in care [16]. More examples of such interventions are needed in this era of “test and treat.”

With funding from the Doris Duke Charitable Foundation’s Operations Research on AIDS Care and Treatment in Africa program, between 2009 and 2013, we developed and operated an intervention, Mobile Antiretroviral Therapy and HIV care (MAP-HC) in rural southwestern Uganda. Our intervention aimed to reduce travel distance and patient cost associated with travel to collect medicines and, indirectly, reduce wait time and crowding at the HIV clinic. We also hypothesized that MAP-HC would improve ART adherence and rates of viral load suppression. We report the results of our program evaluation.

## Methodology

### Study setting

The study was conducted at two sites, Kitagata and Itojo Hospitals, and these are public health facilities located in rural southwestern Uganda. They were selected because they were among the first district-level ART clinics serving a rural population. Kitagata Hospital is located in Sheema District, Uganda, which is predominantly a rural population of subsistence farmers. The area has a difficult terrain and transportation around the district is mainly by motorcycle taxis or *boda bodas*. Itojo Hospital is located just south in Ntungamo District with mixed subsistence farming and livestock as the major sources of livelihood. At both hospitals, majority of patients travel several hours to the hospital to receive their monthly ART refills. By the beginning of 2009, the cumulative number of patients receiving ART was over 520 for Kitagata Hospital and 632 for Itojo Hospital.

### Development and Implementation of MAP-HC

The implementation of the MAP-HC was done in three phases, the Preparation, Identification, and Implementation phases. We engaged health workers and patients at both ART clinics to ensure the process was participatory, but began the development of MAP-HC at Kitagata Hospital where we had already established working relationships with healthcare workers and hospital administration. Once MAP-HC was developed and operational at Kitagata, we repeated our implementation steps at Itojo Hospital.

In the *preparation phase*, which began in March 2009 at Kitagata Hospital and October 2009 at Itojo Hospital, our team began to sensitize ART clients about the intervention. We conducted a baseline survey at the hospital to measure patient demographics, place of residence, distance travelled to the clinic, pre-intervention adherence to ART and viral load suppression. We also defined the catchment area of the clinic and mapped patients’ places of residence by parish and village. We used this information to identify the clusters or zones where majority of patients resided. The catchment area of residence was then divided into 4 zones based on clustering of the patient population. Within each zone, we identified a county- or sub-county-(Health Centre III) or parish-level (Health Centre II) healthcare facilities, which was closer in distance than the district-level hospital (Health Centre IV) to the majority of patients in that zone, to serve as a distribution point to dispense antiretroviral therapy.

In the *identification phase*, clinicians at the hospital interacted with patients and informed them about our proposed intervention. At Kitagata Hospital, where we first began to work, patients were asked to choose whether they would wish to receive their refills at our proposed ART dispensing sites or continue receiving their medications at the hospital. Patients were enthusiastic for our proposed intervention, but expressed a desire for healthcare services in addition to ART refills. We then worked with hospital administration to determine the feasibility and logistics of adding healthcare delivery elements to our intervention. Ultimately, we revised our intervention from being ART dispensing alone (MAP) to one that also provides basic healthcare services (MAP-HC), which is described further below.

Patients were offered MAP-HC services if they had been taking ART for at least 6 months, were considered stable on treatment by the physician, and lived further than five kilometers from the district hospital, a distance determined by the Ministry of Health as being excessive when traveling for one’s healthcare. [17] In making their decision, patients were asked to consider other issues such as privacy and whether the MAP-HC site would be convenient to them.

In the *implementation phase*, we reviewed the appointment dates for all patients that consented to participate in MAP-HC and synchronized appointment and ART refill dates by zone. We obtained buy in from the district-level health service team, local government and the healthcare facility where MAP-HC would be stationed. HIV care providers selected a non-HIV clinic day during the week to operate MAP-HC and chose a day that was also typically lighter at the hospital to allow the team be away.

### MAP-HC intervention

The MAP-HC team consisted of five members: two nurses trained in dispensing ART and monitoring adverse events, a trained medication dispenser, an ART adherence counselor, a driver. The mobile team provided ART refills, adherence counseling, and treatment of concomitant illnesses such as malaria, respiratory tract infections and other conditions that did not require an admission. The physician who ordinarily runs the HIV clinic was required to remain back at the hospital to serve other non-HIV care seeking patients.

The MAP-HC team used the hospital’s own vehicle and the research project investigators provided reimbursement for fuel expenses and field day allowance at government approved rates for health care staff. Patients received ART and basic care at MAP-HC sites for three consecutive visits, but were required to return to the district hospital for the fourth visit in order to be seen by the physician.

#### Data collection

We compared measurements before and after the MAP. First, we conducted a baseline assessment at the two district hospitals to measure baseline features such as waiting time, adherence and viral load. Next, we implemented the project, and then collected post intervention measurements. We measured waiting time at the hospitals and sampled patients for viral load and adherence measurement at the MAP sites. We repeated measurements of patient waiting time, adherence and viral load 12 months after the MAP was implemented.

#### Study outcomes

The primary outcome for this evaluation was viral load suppression. The secondary outcomes were attendance at MAP sites, adherence to antiretroviral medication, and waiting time at the hospital. We hypothesized that adherence and viral load suppression would improve among patients attending MAP and that MAP would decongest the ART clinic at the hospital resulting in reduced waiting time.

To measure patient waiting time at the hospital, we positioned 4 research assistants at the ART Clinic following the order of flow at the clinic. The research assistants were positioned at the entrance/registration area, adherence officer room, clinician consultation rooms, and dispensary. Patients followed this natural order and collected their medications at the dispensary before departure. On arrival at the entrance of the ART clinic, the research assistant wrote the arrival time on a small script of paper, which they handed to the patient. The research assistant at the next stationed wrote the time when the patient completed the visit at each station, and the final research assistant at the dispensary wrote the time and collected the script from the patient before their departure. The typical layout for the clinic at Kitagata Hospital is shown in Figure 1 below.

**Figure 1:**
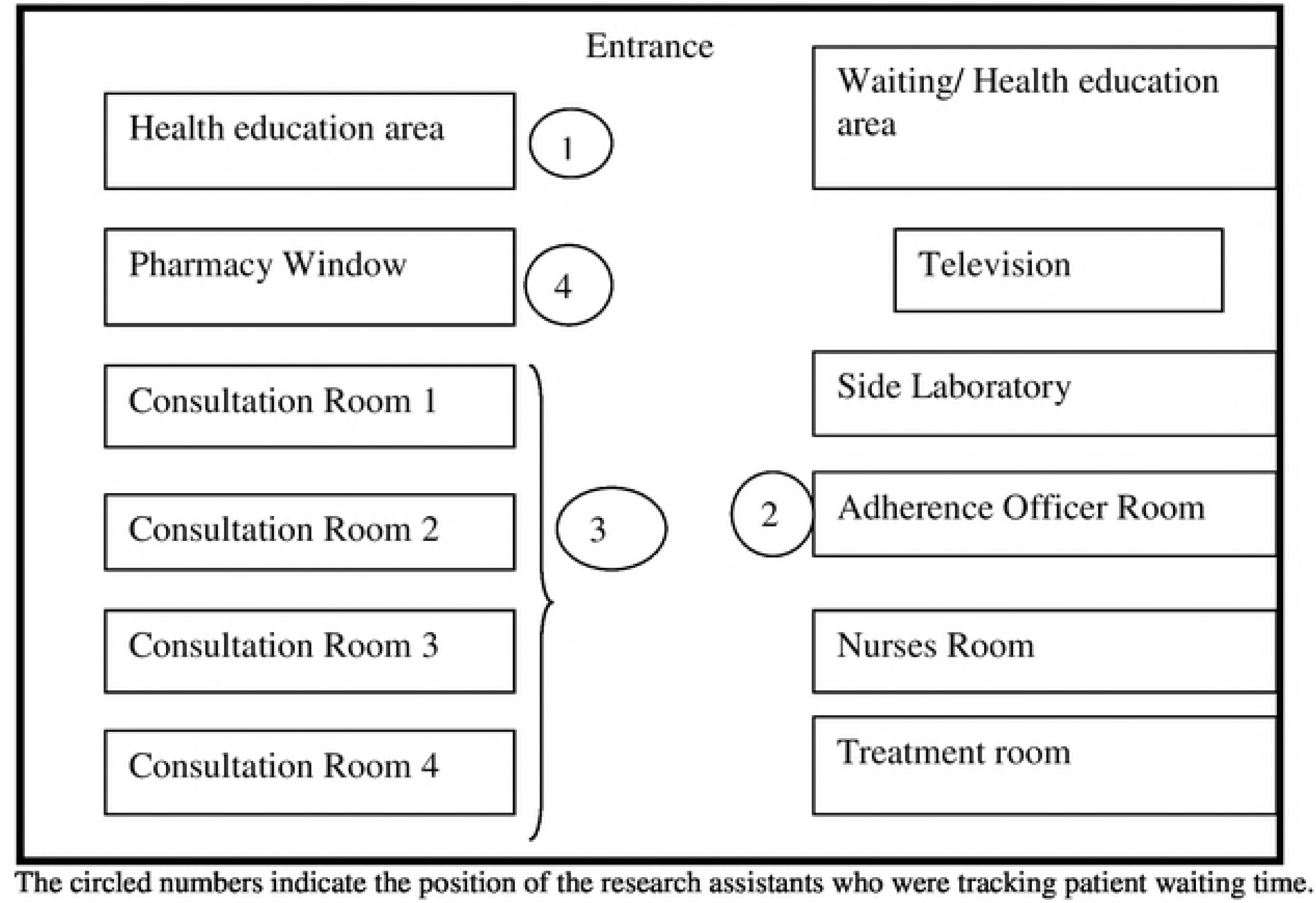
Lay out for the ART clinic at kitagata Hospital and location of research assistants to measure patient waiting time

The circled numbers in Figure 1 represent the positions where the research assistants were positioned to collect the waiting time data.

Adherence was measured using self-report. Blood samples were drawn from the patients and sent to a regional testing laboratory in Mbarara for viral load testing. VL were measured using Amplicor system^®^ from Roche. Results were relayed back to the health care providers 4 weeks after sample collection.

#### Sampling, sample size and data analysis

We used consecutive sampling to collect baseline data. We aimed to interview 300 study participants at baseline to determine the proportion of participants that were living outside of 5km from the hospital. At the follow up for evaluation, we did not calculate sample size. The number was determined by the resources available to conduct viral load testing, as this was the primary outcome for the evaluation.

We summarized the baseline characteristics using 5km as the cut off. This is because the Ministry of Health recommends that persons should reside within a radius of 5km from a health facility [18]. We compared continuous baseline variables using non-parametric tests and categorical outcomes using Chi square tests. We compared the pre- and post-intervention proportions of patients who were adherence or had viral load suppression using Chi square test and a t-test to compare waiting time. Paired tests were not used since the sampling process did not necessarily include the same patients before and after the intervention.

#### Human subjects’ issues

The proposal was approved by the Research Ethics Committee at Mbarara University of Science and Technology and by the Uganda National Council of Science and Technology. Study procedures were explained to the clients at the HIV clinic. Individual written informed consent was obtained and all study participants signed a consent form once they understood study procedures and accepted to participate. Participants were made aware they were free to decline participation in the MAP but continue to receive their medications at the hospital. Confidentiality was maintained by using number identifications for the patients on study materials and questionnaires.

## Results

We conducted baseline assessment interviews among 292 patients at both Kitagata and Itojo Hospital. Almost two thirds were women, with a median age of 37 years. Majority lived more than 5km away from the hospital and the results are shown in Table 1. As expected, those who lived nearer were more likely to walk compared to those who lived further. Participants who lived further than 5 km also had significantly lowed median monthly incomes compared to those who lived near the health facility (p=0.021). Those who lived further than 5km also spent more time and money in travel to the hospital.

**Table 1:** Baseline characteristics of rural patients attending ART clinic at Kitagata and Itojo Hospital, south western Uganda, 2009

| Characteristic | Overall n=292 | Less than 5km<br>n=79 | More than 5km<br>n=213 | p value |
| --- | --- | --- | --- | --- |
| Gender |  |  |  |  |
| Male | 104 (36%) | 39 (49%) | 65 (31%) | 0.003 |
| Female | 188 (64%) | 40 (51%) | 148 (69%) |  |
| Age (median, IQR) | 37 (32, 43) | 36 (32, 42) | 38 (33, 44) | 0.041 |
| Has job income | 183 (63%) | 54 (68%) | 129 (61%) | 0.221 |
| Monthly income USD (median, IQR) | 15 (10, 50) | 25 (12.5, 62) | 15 (10, 45) | 0.021 |
| Transport to clinic: |  |  |  |  |
| Walking | 91 (31%) | 44 (56%) | 47 (22%) | <0.001 |
| Rode a bicycle | 20 (7%) | 6 (8%) | 14 (7%) |  |
| Took a bodaboda | 88 (30%) | 24 (30%) | 64 (30%) |  |
| Took a minibus | 93 (32%) | 5 (6%) | 88 (41%) |  |
| Travel time to clinic (minutes median, IQR) | 90 (60, 140) | 60 (30, 90) | 120 (100, 150) | <0.001 |
| Travel cost USD (median, IQR) | 2.5 (1.5, 3.0) | 1.5 (1.0, 2.0) | 3.0 (2.0, 4.0) | <0.001 |

Participants had comparable 3 and 30 day self-report adherence regardless of whether they lived within or outside 5km from the hospital (p=0.577) and these results are shown in Table 2 below. However, participants who lived further than 5km were more likely to report having ever missed a pill compared to those who lived near (p=0.013). The participants that lived further than 5km were also more likely to report distance as a barrier to their travel to the clinic.

**Table 2:** Characteristics related to antiretroviral treatment among rural patients at Kitagata Hospital, southwestern Uganda

| Characteristic | Overall<br>n=292 | Less than 5km<br>n=79 | More than 5km<br>n=213 | p value |
| --- | --- | --- | --- | --- |
| Months on ART<br>(median, IQR) | 25 (14, 36) | 29 (14, 39) | 24 (14, 36) | 0.57 |
| Missed appointment | 47 (16%) | 9 (11%) | 38 (18%) | 0.134 |
| Distance affects<br>coming to clinic: |  |  |  |  |
| All of the time | 64 (22%) | 5 (6%) | 59 (28%) | <0.001 |
| Most of the time | 44 (15%) | 6 (8%) | 38 (18%) |  |
| Rarely | 112 (38%) | 29 (37%) | 83 (39%) |  |
| Never | 72 (25%) | 39 (49%) | 33 (15%) |  |
| Missed ART dose: |  |  |  |  |
| In last 3 days | 4 (1%) | 0 (0%) | 4 (2%) | 0.577 |
| In last 30 days | 57 (20%) | 16 (20%) | 41 (19%) | 0.847 |
| Missed an entire day<br>of ART in last 30 days | 35 (12%) | 8 (10%) | 27 (13%) | 0.551 |
| Ever missed ART dose | 123 (42%) | 24 (30%) | 99 (45%) | 0.013 |

### Post intervention results

Figure 2 shows a bar graph with attendance at the MAP, with numbers of participants expected at each month visit versus those who actually turned up. The graph shows that consistently, the attendance was nearly 100% and in some instances, exceeded the expected numbers.

**Figure 2:**
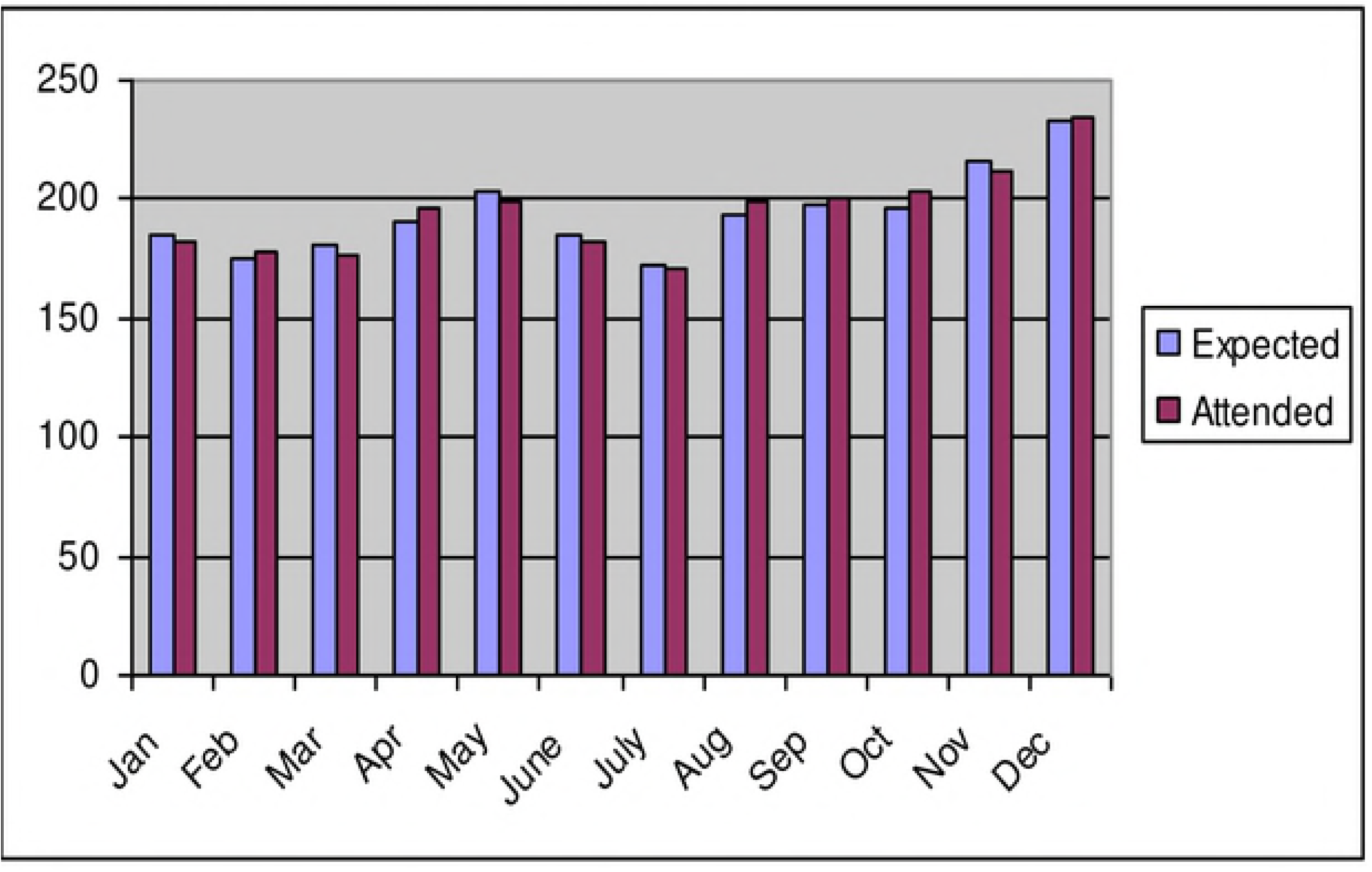
Number of patients attending at the Mobile Antiretrovial therapy program, southwastern Uganda, January - December 2010

The mean waiting time at the hospital was 4.48 hours before the intervention and increased to 4.76 hours after the intervention, however this change was not statistically significant (p=0.13) and these results are shown in Table 3. The proportion of patients who missed an ART dose in the last 30 days dropped from 20% at baseline to 8.5% at 12 months after the intervention and this drop was statistically significant (p=0.009). Similarly, the proportion of patients with detectable viral load significantly dropped from 19.9% to 7.4% after the intervention (p=0.001).

**Table 3:**
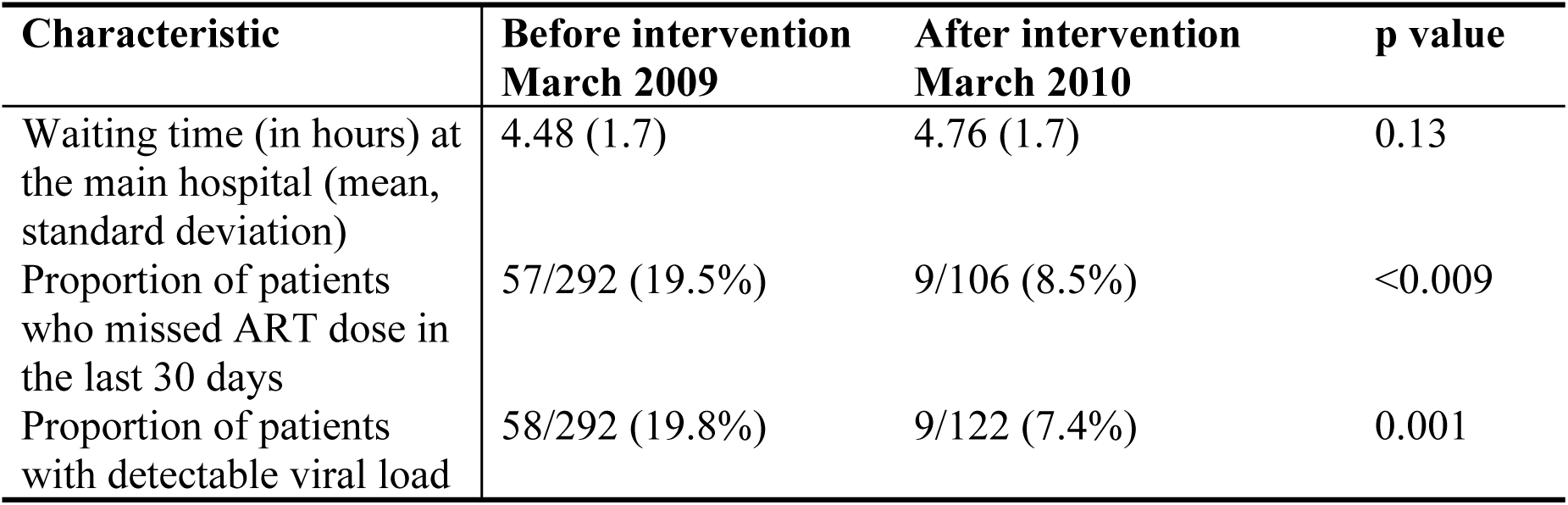
Changes in the treatment outcomes before and after implementation of the Mobile antiretroviral therapy program, southwestern Uganda

## Discussion

In this mobile pharmacy intervention for ART in rural western Uganda, there was significant reduction in the proportion of patients who missed an ART dose in the last 30 days and a significant decrease in the proportion of patients with detectable viral load. However, the intervention had no impact on the waiting time at the district hospital.

Our intervention targeted participants that lived more than 5km from the hospital and our data indicate this was the right target population. Not only did they take more time to travel to the hospital, they also paid more to reach the facility and reported more difficulty trying to reach it. Yet, these participants who lived further away were more likely to earn less per month compared to those who lived nearer, placing more burden on their meager resources to meet their transportation fee to the hospital per month. Those who lived further than 5km were more likely to reside in more rural remote establishments. Although majority of the districts are predominantly rural, the hospital was located in a semi-urban setting. This more rural population needed the intervention.

The hospital ART clinics were overcrowded pre-intervention. We hypothesized that MAP would decongest them by taking away a large proportion who needed refills. Despite the intervention, there was no significant decrease in waiting time at the hospital. On closer scrutiny, the lack of decrease in waiting time came as no surprise because the overall ART clinic population for instance at Kitagata grew from 500 patients to almost 1000 patients within a year following the implementation of the MAP. The potential impact of the intervention was washed away by the exponential growth in the clinic size. This is an important challenge to anticipate as clinic populations are expected to grow even more as Uganda is now implementing the ‘test and treat’ approach to achieve 90-90-90 UNAIDS targets. As efforts to identify and treat more people grow, the burden on health centers grows, and without other approaches to providing treatment, the increased burden could eventually stall or reverse gains made.

Our results agree with other studies in sub Saharan Africa and elsewhere where results show that programs to get patients to receive treatment at lower level health facilities are successful [19] or even have better treatment outcomes compared to those at higher level facilities [20, 21]. Our study did not perform a direct comparison between higher and lower level facilities but instead compared a pre-intervention (district hospital) versus a post-intervention (lower level facility) outcome outcomes. In our evaluation, the better outcomes following the MAP intervention may be attributed to the elimination of distance, a structural barrier to adherence. The results agree with our hypothesis that removal of distance barrier would result in improvement in adherence and viral load suppression. In the United States and other resource rich countries, a related kind of intervention designed to improve adherence and involved the referral of patients who were stable on ART to HIV-focused community pharmacies [22, 23] also showed significant success.

Our results and others elsewhere, suggest that ART programs should consider scale up of ART to proximal health facilities or community pharmacies in rural areas and where patients have to travel long distances and or difficult terrain such as that here in rural western Uganda where our study was based. However, there may be concerns about the long term sustainability of this kind of program, especially where additional financial support is required as was the case in our program. To ensure sustainability, the mobile pharmacies should embrace the concept of decentralization of HIV services and based on a recent review [24] should be considered for integration with other well-known successful interventions such as short messaging services, adherence clubs and peer counseling, or even merge with other outreach programs such as vaccinations.

Our evaluation has important strengths. First, the evaluation was conducted in a rural population. This is important because majority of populations in sub Saharan Africa and Uganda live in rural areas but also because patients in rural areas are more likely to experience distance as a barrier to health care seeking. Second, we measured several markers of impact of the MAP including crowding, adherence and viral load suppression. Although self-reported adherence may be influenced by social desirability bias, viral load suppression is insulated from subjective reporting.

Our study has some limitations. In our sampling, some of the patients involved in the pre-intervention assessment were not necessarily involved in the post intervention VL and adherence measurement. In the post-intervention assessment, we randomly sampled MAP patients for viral load measurement, regardless of whether they had been involved in the baseline assessment or not. Although this sampling approach partly engages components of convenience in the sampling, it is unlikely to have influenced the results because all patients receiving care at a MAP site lived more than 5 km from the hospital and reported distance as a barrier to adherence. The sampling approach is consistent with other pseudo-experimental study designs that involve a pre- and post-evaluation comparison. Lastly, we did not conduct a cost-effectiveness analysis for this intervention and future evaluations should incorporate data collection to support this analysis.

In conclusion, our study has showed that a mobile pharmacy intervention in rural Uganda is feasible and resulted in improvement in adherence and viral load suppression. Although it did not reduce patient waiting time at the clinic, we recommend a scale-up of this intervention in rural areas where patients face challenges of transportation to the clinic.

## Acknowledgements

We would like to thank the staff and administration of Kitagata and Itojo Hospitals in western Uganda for supporting our research team in the recruitment and data collection process. We thank the Bushenyi district local government for supporting the implementation of the mobile ART program, and last but not least the patients who participated in the implementation and evaluation of the program. This work was funded by Oerations Research on AIDS Care and Treatment in Africa (ORACTA) Award (2007012) from the Doris Duke Charitable Foundation.

## References

1. Gross R, Bilker WB, Friedman HM, Strom BL. Effect of adherence to newly initiated antiretroviral therapy on plasma viral load. AIDS (London, England). 2001;15(16):2109–17. Epub 2001/10/31. PubMed PMID: 11684930.

2. Bangsberg DR, Hecht FM, Charlebois ED, Zolopa AR, Holodniy M, Sheiner L, et al. Adherence to protease inhibitors, HIV-1 viral load, and development of drug resistance in an indigent population. AIDS (London, England). 2000;14(4):357–66. Epub 2000/04/19. PubMed PMID: 10770537.

3. Arnsten JH, Demas PA, Grant RW, Gourevitch MN, Farzadegan H, Howard AA, et al. Impact of active drug use on antiretroviral therapy adherence and viral suppression in HIV-infected drug users. Journal of general internal medicine. 2002;17(5):377–81. Epub 2002/06/06. PubMed PMID: 12047736; PubMed Central PMCID: PMCPMC1495042.

4. Sethi AK, Celentano DD, Gange SJ, Moore RD, Gallant JE. Association between adherence to antiretroviral therapy and human immunodeficiency virus drug resistance. Clinical infectious diseases : an official publication of the Infectious Diseases Society of America. 2003;37(8):1112–8. Epub 2003/10/03. doi: 10.1086/378301. PubMed PMID: 14523777.

5. Mills EJ, Nachega JB, Buchan I, Orbinski J, Attaran A, Singh S, et al. Adherence to antiretroviral therapy in sub-Saharan Africa and North America: a meta-analysis. Jama. 2006;296(6):679–90. Epub 2006/08/10. doi: 10.1001/jama.296.6.679. PubMed PMID: 16896111.

6. Shubber Z, Mills EJ, Nachega JB, Vreeman R, Freitas M, Bock P, et al. Patient-Reported Barriers to Adherence to Antiretroviral Therapy: A Systematic Review and Meta-Analysis. PLoS medicine. 2016;13(11):e1002183. Epub 2016/11/30. doi: 10.1371/journal.pmed.1002183. PubMed PMID: 27898679; PubMed Central PMCID: PMCPMC5127502.

7. Bajunirwe F, Arts EJ, Tisch DJ, King CH, Debanne SM, Sethi AK. Adherence and treatment response among HIV-1-infected adults receiving antiretroviral therapy in a rural government hospital in Southwestern Uganda. Journal of the International Association of Physicians in AIDS Care (Chicago, Ill : 2002). 2009;8(2):139–47. Epub 2009/03/05. doi: 10.1177/1545109709332470. PubMed PMID: 19258526.

8. Akullian AN, Mukose A, Levine GA, Babigumira JB. People living with HIV travel farther to access healthcare: a population-based geographic analysis from rural Uganda. Journal of the International AIDS Society. 2016;19(1):20171. Epub 2016/02/13. doi: 10.7448/ias.19.1.20171. PubMed PMID: 26869359; PubMed Central PMCID: PMCPMC4751409.

9. Lankowski AJ, Siedner MJ, Bangsberg DR, Tsai AC. Impact of geographic and transportation-related barriers on HIV outcomes in sub-Saharan Africa: a systematic review. AIDS and behavior. 2014;18(7):1199–223. Epub 2014/02/25. doi: 10.1007/s10461-014-0729-8. PubMed PMID: 24563115; PubMed Central PMCID: PMCPMC4047127.

10. Tuller DM, Bangsberg DR, Senkungu J, Ware NC, Emenyonu N, Weiser SD. Transportation costs impede sustained adherence and access to HAART in a clinic population in southwestern Uganda: a qualitative study. AIDS and behavior. 2010;14(4):778–84. Epub 2009/03/14. doi: 10.1007/s10461-009-9533-2. PubMed PMID: 19283464; PubMed Central PMCID: PMCPMC2888948.

11. Biadgilign S, Deribew A, Amberbir A, Deribe K. Barriers and facilitators to antiretroviral medication adherence among HIV-infected paediatric patients in Ethiopia: A qualitative study. SAHARA J : journal of Social Aspects of HIV/AIDS Research Alliance. 2009;6(4):148–54. Epub 2010/05/21. PubMed PMID: 20485854.

12. Hardon AP, Akurut D, Comoro C, Ekezie C, Irunde HF, Gerrits T, et al. Hunger, waiting time and transport costs: time to confront challenges to ART adherence in Africa. AIDS care. 2007;19(5):658–65. Epub 2007/05/17. doi: 10.1080/09540120701244943. PubMed PMID: 17505927.

13. Consolidated guidelines on the use of antiretroviral drugs for treating and preventing HIV infection Recommendations for a public health approach -Second edition 2016 [March 30, 2018]. http://www.who.int/hiv/pub/arv/arv-2016/en/].

14. Bemelmans M, van den Akker T, Ford N, Philips M, Zachariah R, Harries A, et al. Providing universal access to antiretroviral therapy in Thyolo, Malawi through task shifting and decentralization of HIV/AIDS care. Tropical medicine & international health: TM & IH. 2010;15(12):1413–20. Epub 2010/10/21. doi: 10.1111/j.1365-3156.2010.02649.x. PubMed PMID: 20958897.

15. Bedelu M, Ford N, Hilderbrand K, Reuter H. Implementing antiretroviral therapy in rural communities: the Lusikisiki model of decentralized HIV/AIDS care. The Journal of infectious diseases. 2007;196 Suppl 3:S464–8. Epub 2008/01/10. doi: 10.1086/521114. PubMed PMID: 18181695.

16. Avong YK, Aliyu GG, Jatau B, Gurumnaan R, Danat N, Kayode GA, et al. Integrating community pharmacy into community based anti-retroviral therapy program: A pilot implementation in Abuja, Nigeria. PLoS One. 2018;13(1):e0190286. Epub 2018/01/11. doi: 10.1371/journal.pone.0190286. PubMed PMID: 29320531; PubMed Central PMCID: PMCPMC5761864.

17. Guidelines for Designation, Establishment and Upgrading of Health Facilities in Uganda; http://health.go.ug/docs/guidelines.pdf Accessed December 10, 2018. 2011.

18. HEALTH SECTOR DEVELOPMENT PLAN 2015/16 - 2019/20: Ministry of Health, Uganda. Available from: http://health.go.ug/sites/default/files/Health%20Sector%20Development%20Plan%202015-16_2019-20.pdf

19. Grimsrud A, Sharp J, Kalombo C, Bekker LG, Myer L. Implementation of community-based adherence clubs for stable antiretroviral therapy patients in Cape Town, South Africa. Journal of the International AIDS Society. 2015;18:19984. Epub 2015/05/30. doi: 10.7448/ias.18.1.19984. PubMed PMID: 26022654; PubMed Central PMCID: PMCPMC4444752.

20. Fatti G, Grimwood A, Bock P. Better antiretroviral therapy outcomes at primary healthcare facilities: an evaluation of three tiers of ART services in four South African provinces. PLoS One. 2010;5(9):e12888. Epub 2010/09/30. doi: 10.1371/journal.pone.0012888. PubMed PMID: 20877631; PubMed Central PMCID: PMCPMC2943483.

21. Grimsrud A, Lesosky M, Kalombo C, Bekker LG, Myer L. Implementation and Operational Research: Community-Based Adherence Clubs for the Management of Stable Antiretroviral Therapy Patients in Cape Town, South Africa: A Cohort Study. Journal of acquired immune deficiency syndromes (1999). 2016;71(1):e16–23. Epub 2015/10/17. doi: 10.1097/qai.0000000000000863. PubMed PMID: 26473798.

22. Cocohoba JM, Murphy P, Pietrandoni G, Guglielmo BJ. Improved antiretroviral refill adherence in HIV-focused community pharmacies. Journal of the American Pharmacists Association : JAPhA. 2012;52(5):e67–73. Epub 2012/10/02. doi: 10.1331/JAPhA.2012.11112. PubMed PMID: 23023860; PubMed Central PMCID: PMCPMC4607273.

23. Hirsch JD, Rosenquist A, Best BM, Miller TA, Gilmer TP. Evaluation of the first year of a pilot program in community pharmacy: HIV/AIDS medication therapy management for Medi-Cal beneficiaries. Journal of managed care pharmacy : JMCP. 2009;15(1):32–41. Epub 2009/01/08. doi: 10.18553/jmcp.2009.15.1.32. PubMed PMID: 19125548.

24. Haberer JE, Sabin L, Amico KR, Orrell C, Galarraga O, Tsai AC, et al. Improving antiretroviral therapy adherence in resource-limited settings at scale: a discussion of interventions and recommendations. Journal of the International AIDS Society. 2017;20(1):21371. Epub 2017/06/21. doi: 10.7448/ias.20.1.21371. PubMed PMID: 28630651; PubMed Central PMCID: PMCPMC5467606.

